# Improved protein structure prediction using predicted inter-residue orientations

**DOI:** 10.1101/846279

**Authors:** Jianyi Yang, Ivan Anishchenko, Hahnbeom Park, Zhenling Peng, Sergey Ovchinnikov, David Baker

## Abstract

The prediction of inter-residue contacts and distances from co-evolutionary data using deep learning has considerably advanced protein structure prediction. Here we build on these advances by developing a deep residual network for predicting inter-residue orientations in addition to distances, and a Rosetta constrained energy minimization protocol for rapidly and accurately generating structure models guided by these restraints. In benchmark tests on CASP13 and CAMEO derived sets, the method outperforms all previously described structure prediction methods. Although trained entirely on native proteins, the network consistently assigns higher probability to *de novo* designed proteins, identifying the key fold determining residues and providing an independent quantitative measure of the “ideality” of a protein structure. The method promises to be useful for a broad range of protein structure prediction and design problems.

## Introduction

Clear progress in protein structure prediction was evident in the recent CASP13 structure prediction challenge (1). Multiple groups showed that application of deep learning-based methods to the protein structure prediction problem makes it possible to generate fold-level accuracy models of proteins lacking homologs in the Protein Data Bank (PDB) (2) directly from multiple sequence alignments (MSAs) (3–6). In particular, AlphaFold (A7D) from DeepMind (7) and Jinbo Xu with RaptorX (4) showed that distances between residues (not just the presence or absence of a contact) could be accurately predicted by deep learning on residue coevolution data. The three top-performing groups (A7D, Zhang-Server and RaptorX) all used deep residual convolutional networks with dilation, with input features of coevolutionary couplings derived from MSAs either using pseudo-likelihood or by covariance matrix inversion. Because these deep learning-based methods produce more complete and accurate predicted distance information, 3D structures can be generated by direct optimization. For example, the Xu group (4) used CNS (8) and the AlphaFold group (7) used gradient-descent following conversion of the predicted distances into smooth restraints. Progress was also evident in protein structure refinement at CASP13 using energy guided refinement (9–11).

In this work, we integrate and build upon the CASP13 advances. We show that through extension of deep learning-based prediction to inter-residue orientations in addition to distances, and the development of a Rosetta-based optimization method that supplements the predicted restraints with components of the Rosetta energy function, still more accurate models can be generated. We also explore applications of the model to the protein design problem. To facilitate further development in this rapidly moving field, we make all the codes for the improved method available.

## Results and Discussion

### Overview of the method

The key components of our method (named trRosetta, transform-restrained Rosetta) include: (i) a deep residual convolutional network which takes an MSA as the input and outputs information on the relative distances and orientations of all residue pairs in the protein, and (ii) a fast Rosetta model building protocol based on restrained minimization with distance and orientation restraints derived from the network outputs.

#### (i) Predicting inter-residue geometries from MSAs using a deep neural network

Unlike most other approaches to contact/distance predictions from MSAs, in addition to C_*β*_-C_*β*_ distances, we also sought to predict inter-residue orientations (Fig. 1A). Orientations between residues 1 and 2 are represented by 3 dihedral (*ω, θ*_12_, *θ*_21_) and 2 planar angles (*φ*_12_, *φ*_21_) as shown on Fig. 1A. The *ω* dihedral measures rotation along the virtual axis connecting the C_*β*_ atoms of the two residues, and *θ*_12_, *φ*_12_ (*θ*_21_, *φ*_21_) angles define the direction in which the C_*β*_ atom of residue 2 (1) is seen from residue 1 (2). Unlike *d* and *ω, θ* and *φ* coordinates are asymmetric and depend on the order of residues (1-2 and 2-1 pairs yield different coordinates which is the reason why *θ* and *φ* maps on Fig. S1 are asymmetric). Together, the 6 parameters *d, ω, θ*_12_, *φ*_12_, *θ*_21_, *φ*_21_ fully define the relative positions of the backbone atoms of two residues. All the coordinates show characteristic patterns (Fig. S1), and we hypothesized that a deep neural network could be trained to predict these.

**Fig. 1:**
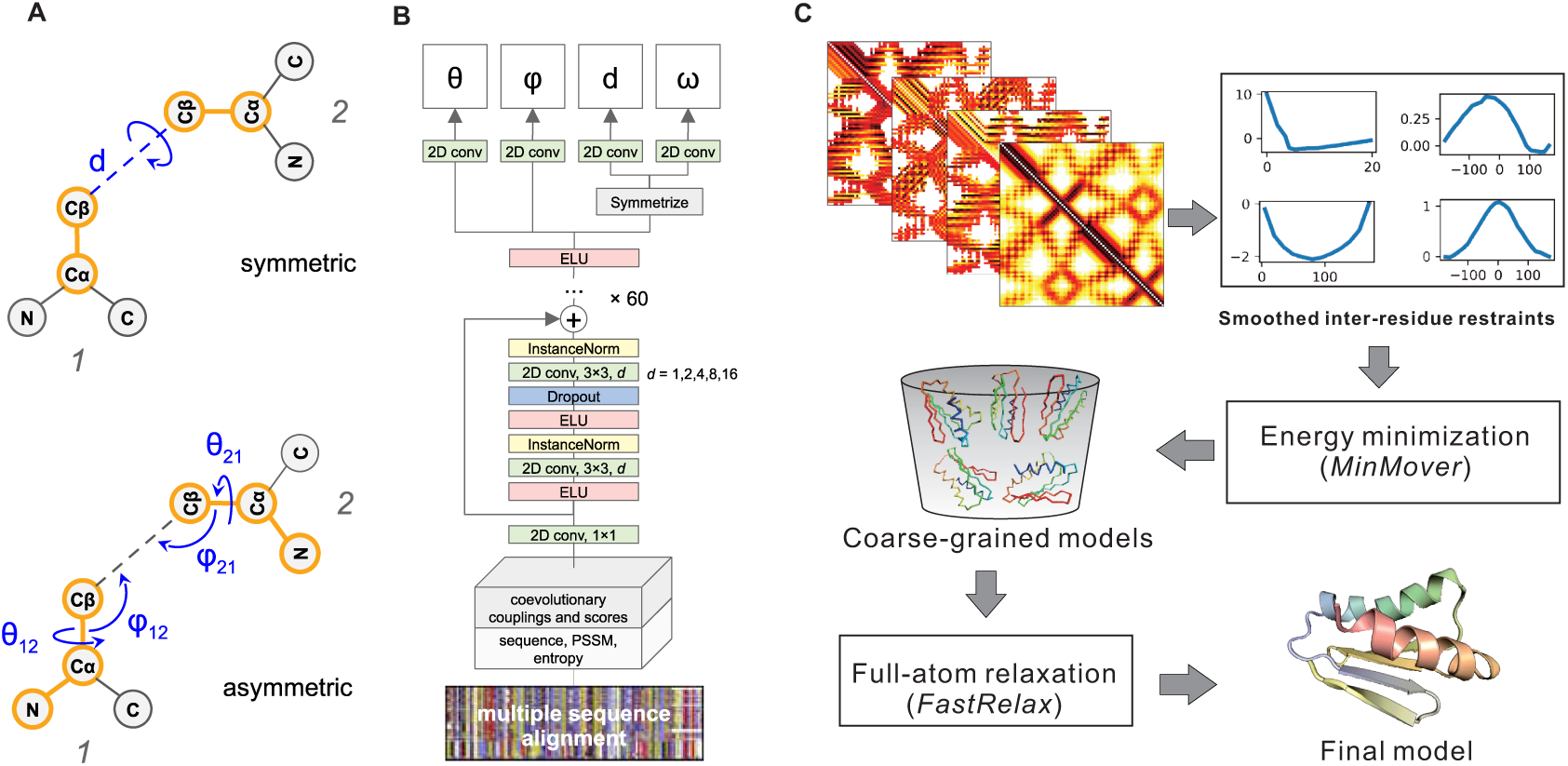
Predicting inter-residue geometries and protein 3D structure from a multiple sequence alignment. (A) Representation of the rigid body transform from one residue to another using angles and distances. (B) Architecture of the deep neural network with multi-objective training to predict inter-residue geometries from an MSA. (C) Outline of the structure modeling protocol based on the restraints derived from the predicted distance and orientation (see Methods for details).

The overall architecture of the network is similar to those recently described for distance and contact prediction (3, 4, 7, 12). Following RaptorX-Contact (4, 12) and AlphaFold (7), we learn probability distributions over distances, and extend this to orientation features. The central part of the network is a stack of dilated residual convolutional blocks which gradually transforms 1- and 2-site features derived from the MSA of the target to predict inter-residue geometries for residue pairs (Fig. 1B) with C_*β*_ atoms closer than 20 Å. The distance range (2-20 Å) is binned into 36 equally spaced segments, 0.5 Å each, plus one bin indicating that residues are not in contact. After the last convolutional layer, the softmax function is applied to estimate the probability for each of these bins. Similarly, *ω, θ* dihedrals and *φ* angle are binned into 24, 24 and 12, respectively, with 15° segments (+ one no-contact bin) and are predicted by separate branches of the network. Branching takes place at the very top of the network, with each branch consisting of a single convolutional layer followed by softmax. The premise for such hard parameter sharing at the downstream layers of the networks is that correlations between the different objectives (i.e. orientations and distance) may be learned by the network, potentially yielding better predictions for the individual features. We used cross-entropy to measure the loss for all branches; the total loss is the sum over the 4 per-branch losses with equal weight. Previous work (4) implicitly captured some orientation information by predicting multiple inter-residue distances (C_β_–C_β_, C_α_– C_α_, C_α_–C_g_, C_g_–C_g_, and N–O), but in contrast to our multi-task learning approach a separate network was used for each of the objectives. Our network was trained on a non-redundant (at 30% sequence identity) dataset from PDB consisting of 15,051 proteins (structure release dates before May 1st, 2018). The trained network is available for download at https://github.com/gjoni/trRosetta.

We couple the derivation of residue-residue couplings from MSAs by covariance matrix inversion to the network by making the former part of the computation graph in TensorFlow (13). Sequence reweighting, calculation of 1-site amino acid frequencies, entropies, and coevolutionary couplings and related scores take place on the GPU, and the extracted features are passed into the convolutional layers of the network (most previous approaches have precomputed these terms). We took advantage of our recent observation (14) that with proper regularization, covariance matrix inversion yields inter-residue couplings (see Methods) with only minor decrease in accuracy compared to pseudo-likelihood approaches like GREMLIN (15) (the latter are prohibitively slow for direct integration into the network). Since the MSA processing steps are now cheap to compute (compared to the forward and backward passes through the network during parameter training), this coupled network architecture allows for data augmentation by massive MSA subsampling during training. At each training epoch, we use a randomly selected subset of sequences from each original MSA, so that each time the network operates on different inputs.

#### (ii) Structure modeling from predicted inter-residue geometries

Following AlphaFold, we generated 3D structures from the predicted distances and orientations using constrained minimization (Fig. 1C). Discrete probability distributions over the predicted orientation and distance bins were converted into inter-residue interaction potentials by normalizing all the probabilities by the corresponding probability at the last bin (see Methods), and smoothing using the *spline* function in Rosetta. These distance and orientation dependent potentials were used as restraints together with the Rosetta centroid level (coarse grained) energy function (16), and folded structures satisfying the restraints were generated starting from conformations with randomly selected backbone dihedral angles by three rounds of quasi-Newton minimization within Rosetta. Only short-range (sequence separation < 12) restraints were included in the first round; medium-range (sequence separation < 24) restraints were added in the second round, and all were included in the third. A total of 150 coarse-grained models were generated using different sets of restraints obtained by selecting different probability thresholds for inclusion of the predicted distances and orientations in modeling.

The 50 lowest-energy backbone+centroid models were then subject to Rosetta full-atom relaxation including the distance and orientation restraints, to add in sidechains and make the structures physically plausible. The lowest-energy full-atom model was then selected as the final model. The structure generation protocol is implemented in PyRosetta (17) and is available as a web server at http://yanglab.nankai.edu.cn/trRosetta/.

### Benchmark tests on CASP13 and CAMEO datasets

#### Accuracy of predicted inter-residue geometries

We tested the performance of our network on 31 FM (free modeling) targets from CASP13 (none of these were included in the training set, which is based on a pre-CASP PDB set). The precision of the derived contacts, defined as the fraction of top *L*/*n* (*n* = 1, 2, 5) predicted contacts realized in the native structure, is summarized in Tables 1 and S1. For the highest probability 7.5% of the distance/orientation predictions (Fig. 2C), there is a good correlation between modes of the predicted distance/orientation distributions and the observed values (Fig. 2C): Pearson’s *r* for distances is 0.72, and circular correlation *r*_*c*_ (18) for *ω, θ* and *φ* are 0.62, 0.77, 0.60, respectively. The predicted probability of the top *L* long+medium-range contacts correlates well (*r* = 0.84) with their actual precision (Fig. 2B). This correlation between predicted probability and actual precision allows us to further improve the results by feeding a variety of MSAs generated with different *e*-value cutoffs or originating from searches against different databases into the network and picking the one which generates predictions with the highest predicted accuracy.

**Table 1.** The precision (%) of the top *L* derived contacts on the 31 CASP13 FM domains and the CAMEO targets compared with the top two contact predictors from CASP13. The values for other methods are slightly different from those listed on the CASP13 website, probably due to different treatment of target length *L* (i.e., length of full sequence or length of domain structures; the latter is used here). The sequence separation between two residues *i* and *j* is denoted by *s* (= |*i*-*j*|).

| Method | CASP13 FM domains |  | CAMEO very hard targets |  |
| --- | --- | --- | --- | --- |
| | $s \geq 24$ | $s \geq 12$ | $s \geq 24$ | $s \geq 12$ |
| RaptorX-Contact | 44.7 | 61.3 | NA | NA |
| TripleRes | 42.3 | 60.9 | NA | NA |
| trRosetta | 51.9 | 70.2 | 48.0 | 62.8 |
| Baseline <sup>a</sup> | 44.3 | 60.7 | 41.6 | 57.5 |
| Baseline+1 <sup>b</sup> | 46.0 | 62.2 | 43.1 | 57.4 |
| Baseline+1+2 <sup>c</sup> | 48.2 | 64.6 | 44.4 | 58.7 |
| Baseline+1+2+3 <sup>d</sup> | 51.3 | 69.3 | 46.1 | 61.4 |

<sup>a</sup> Baseline trRosetta model consists of 36 residual blocks and was trained without MSA subsampling or selection to predict distances only.
<sup>b</sup> 1 - adding MSA subsampling during training
<sup>c</sup> 2 - extending the network to predict orientations
<sup>d</sup> 3 - MSA selection based on predicted probability of the top $L$ long+medium-range contacts

**Fig. 2:**
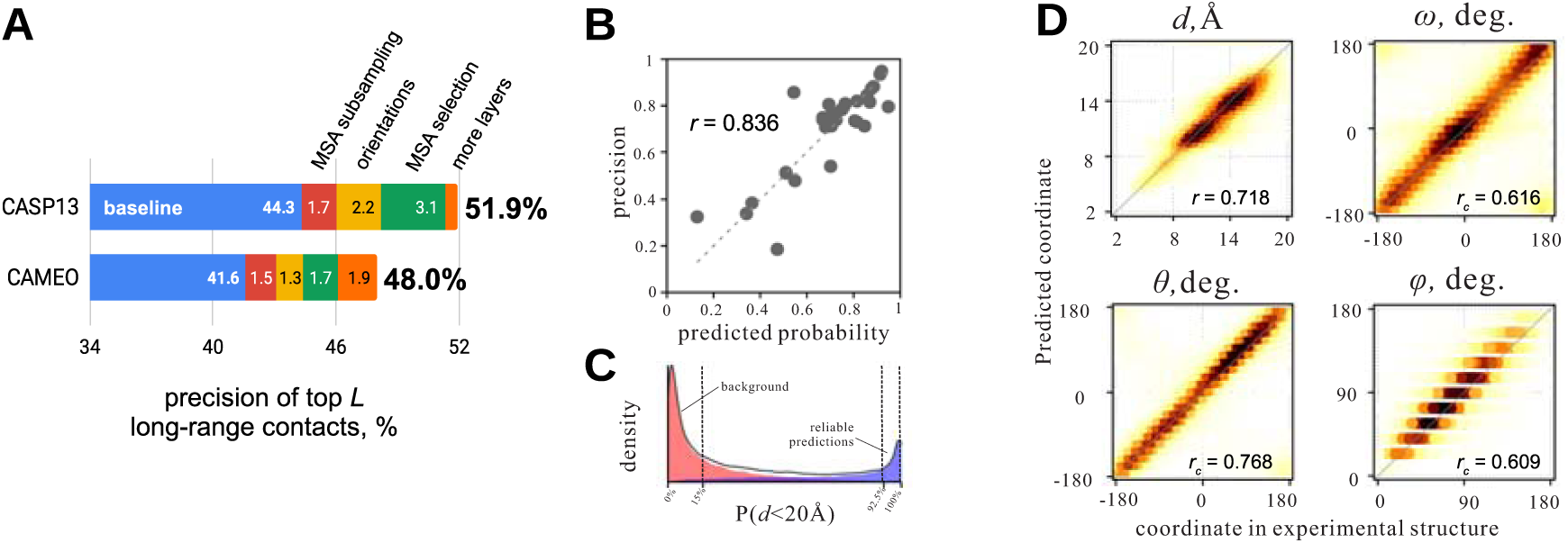
Accuracy of predicted inter-residue geometries. (A) Contribution of different factors to the increase in trRosetta performance on CASP13’s free modeling and CAMEO’s very hard targets. Incorporation of MSA subsampling, orientations and MSA selection in the modeling pipeline increases precision of the top *L* long-range predicted contacts by 1.7% (red bar), 2.2% (yellow) and 3.1% (green) respectively, and increasing the depth of the network from 36 to 61 residual blocks boosts the performance by an additional 0.6% (orange bar). (B) Correlation between predicted probability of the top *L* long+medium-range contacts and their actual precision measured based on the native structures. (C) Distribution of predicted probabilities for residue pairs to be within 20 Å in the native structure; populations in blue and red correspond to residue pairs with *d* ≤ 20 Å and *d* > 20 Å in experimental structures, respectively. Confident predictions are clustered at probability values *P*(*d* < 20 Å) > 92.5%; probabilities for unreliable background predictions are predominantly < 15%. (D) Correlations between actual rigid body transform parameters from the experimental structures with the modes of the predicted distributions for the most reliable long- and medium-range contacts from the top 7.5% percentile; color coding indicates probability density.

#### Comparison with baseline network

We evaluated our extensions to previous approaches by generating a baseline model to predict distances only, with no MSA subsampling and selection; the contact prediction accuracy of this network is comparable to previously described models (3, 12, 19, 20). Incorporating MSA subsampling during training and extending the network to also predict inter-residue orientations improve contact prediction accuracy by 1.7% and 2.2%, respectively. Subsequent alignment selection improves performance an additional 3.1% on the CASP13 FM set (Table 1, last row). The improvements described above, together with increasing the number of layers in the network increase the accuracy of predicted contacts by 7.6% over the baseline network on the CASP13 FM set. Although we ensured that there is no overlap between the training and test sets by selecting pre-CASP PDBs only (before 05/01/2018), our model was trained at a later date when more sequences were available; we also included metagenomic sequence data. Hence, we may be overestimating the gap in performance between our method and those used by other groups in CASP13; future blind tests in CASP will be important in confirming these improvements. Nevertheless, the gain in performance with respect to the baseline model is independent of the possible variations in the training sets and sequence databases. All the targets in the CAMEO validation set below are more recent than both structural and sequence data in the training set.

#### Accuracy of predicted structure models

We tested our method on the CASP13 FM targets, with results shown in Fig. 3. The average TM-score (21) of our method is 0.625, which is 27.3% higher than that (0.491) by the top Server group Zhang-Server (Fig. 3A). Our method also outperforms the top Human group A7D by 6.5% (0.625 vs 0.587; Fig. 3B). The relatively poor performance on T1021s3-D2 (the outlier in the upper triangle of Fig. 3B) reflects the MSA generation procedure: the majority of sequence homologs in the full-length MSA for T1021S3 only cover the first of the two domains; performance is significantly improved (TM-score increased from 0.38 to 0.63; the TM-score of A7D model is 0.672) using a domain-specific MSA. An example of the improved performance of our method is shown in Fig. 3C for the CASP13 target T0950; the TM-score of this model is 0.716, while the highest values obtained during CASP13 are: RaptorX-DeepModeller (0.56), BAKER-ROSETTASERVER (0.46), Zhang-Server (0.44) and A7D (0.43).

**Fig. 3.**
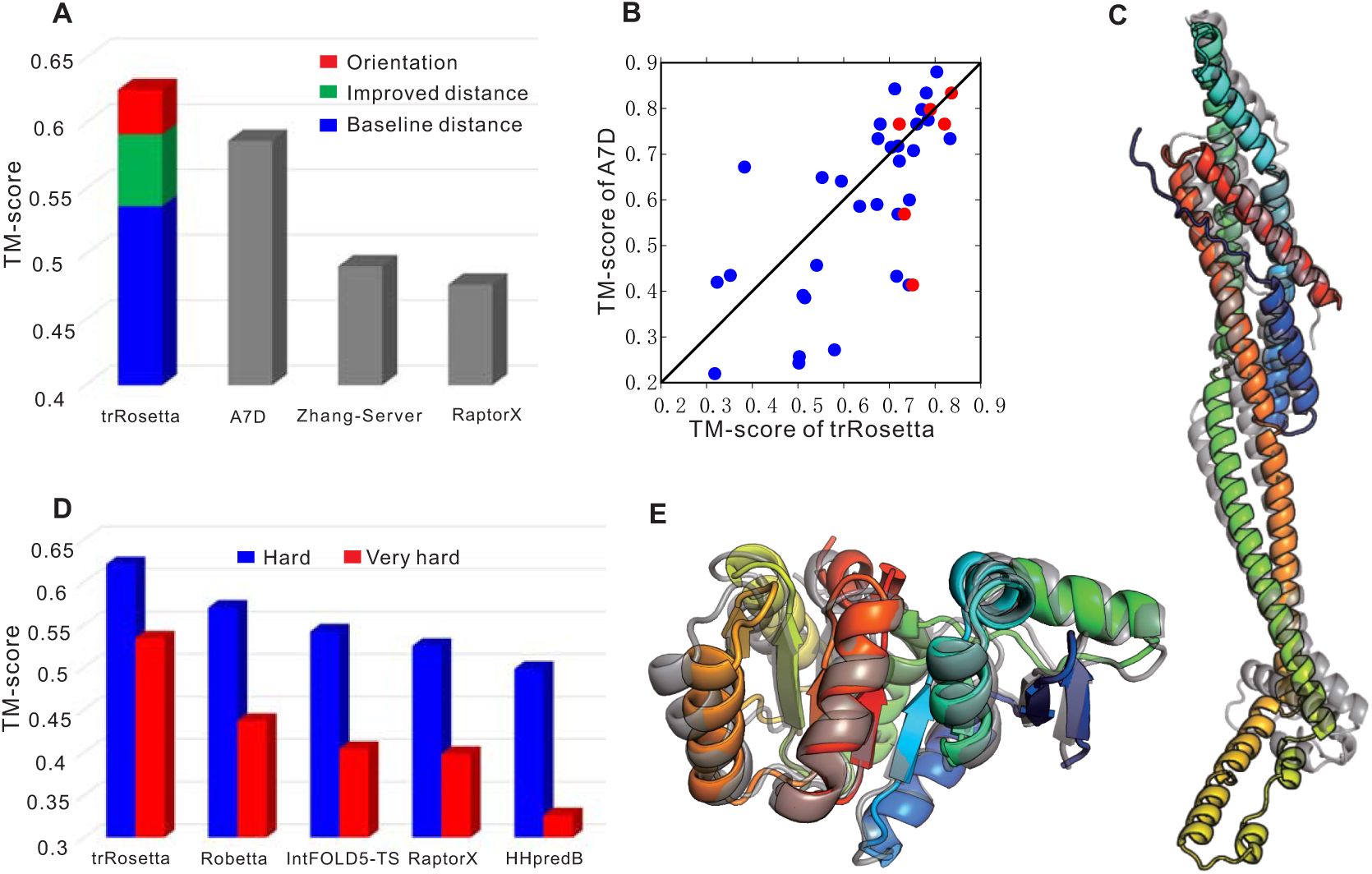
Comparison of model accuracy. (A) average TM-score of all methods on the 31 FM targets of CASP13. The colored stacked bar indicates the contributions of different components to our method. A7D was the top human group in CASP 13; Zhang-Server and RaptorX were the top two server groups. (B) head-to-head comparison between our method and the A7D’s TM-scores over the 31 FM targets (blue points; red points are for six targets with extensive refinement). (C) structures for the CASP13 target T0950; the native structure and the predicted model are shown in gray and rainbow cartoons, respectively. (D) Comparison between our method and the top servers from the CAMEO experiments. (E) Native structure (in gray) and the predicted model (in rainbow) for CAMEO target 5WB4_H. In all of these comparisons, it should be emphasized that the CASP and CAMEO predictions, unlike ours, were made blindly.

Fig. 3A deconstructs the contributions to the improved performance of the different components of our approach. When modeling is only guided by the distance predictions from the baseline network (no orientations and no MSA subsampling and selection; ‘baseline’ bar in Fig. 2A), the TM-score is 0.537, lower than A7D but significantly higher than Zhang-Server and RaptorX. When predicted distances from the complete network are used, the TM-score increases to 0.592, higher than that of A7D. When the orientation distributions are included, the TM-score is further increased to 0.625. The folding is driven by restraints; very similar models are generated without the Rosetta centroid terms, and very poor models are generated without the restraints. To compare our Rosetta minimization protocol (trRosetta) to CNS (8), we obtained predicted distance restraints and structure models for all CASP13 FM targets from the RaptorX-Contacts server (which uses CNS for structure modeling (4)), and used the distance restraints to generate models with trRosetta. The average TM-score of the trRosetta models is 0.45 compared to 0.36 for the RaptorX CNS models; the improvement is likely due to both improved sampling and the supplementation of the distance information with the general Rosetta centroid energy function.

#### Comparison between distance and orientation-based folding

Both predicted distance and orientation can guide folding alone. The average TM-score of coarse-grained models for the CASP13 FM targets is 0.572 when folding with predicted orientation alone and 0.552 when folding with predicted distance only. Detailed head-to-head comparisons are shown in Fig. S2A. After relaxation, models are improved for both. For example, for orientation/distance-based folding, the TM-score increases to 0.58/0.59. Fig. S2B shows that the TM-score difference between the orientation- and distance-based models becomes smaller after relaxation. The differences in model quality for the models generated using either source of information alone suggest that the two are complementary, and indeed better models are generated using both distance and orientation information (Fig. S2).

#### Validation on hard targets from the CAMEO experiments

We further tested our method on 131 hard targets from the CAMEO experiments (22) over the 6 months between 2018.12.08 and 2019.06.01. The results for contact prediction are summarized in Table 1 and Fig. 2A; as in the case of the CASP13 targets, the new method improves over the baseline network. The results for structure modeling are shown in Fig. 3D. The contributions of different components to our method are presented in Fig. S4. On these targets, the average TM-score of our method is 0.621, which is 8.9% and 24.7% higher than Robetta and HHpredB, respectively. We note that the definition of ‘hard’ is looser than the CASP definition; a ‘hard’ target from CAMEO can have close templates in PDB. Making the definition of ‘hard’ more stringent by requiring the TM-score of the HHpredB server to be less than 0.5 reduces the number of targets to 66. On this harder set, the TM-score for our method is 0.534, 22% higher than the top server Robetta and 63.8% higher than the baseline server HHpredB. Fig. 3E gives one example CAMEO target where our method predicts very accurate models (5WB4_H). For this target, the TM-scores of the template-based models by HHpredB, IntFOLD5-TS and RaptorX are about 0.4. In comparison, the TM-score of our predicted model is 0.921, which is also higher than the top server Robetta (0.879).

#### Accuracy estimation for predicted structure models

We sought to predict the TM-score of the final structure model using the 131 hard targets from CAMEO. We found that, unlike direct-coupling based methods such as GREMLIN, the depth of the MSA did not have a good correlation with the accuracy of the derived contacts. Instead, a high correlation (Pearson’s *r* = 0.90) between the average probability of the top predicted contacts and the actual precision was observed (Fig. S3A). The average contact probability also correlates well with the TM-score of the final structure models (*r* = 0.71, Fig. S3B). To obtain a structure-based accuracy metric, we re-relaxed the top 10 models without any restraints. The average pairwise TM-score between these 10 non-constrained models also correlates with the TM-score of the final models (*r* = 0.65, Fig. S3C). Linear regression against the average contact probability and the extent of structural displacement without the restraints gave a quite good correlation between predicted and actual TM-score (*r* = 0.84; Fig. S3D). This method is used to provide an estimated model accuracy.

#### Refinement of predicted models

As noted above, CASP13 showed that protein structure refinement methods can consistently improve models for cases where the sampling problem is more tractable (smaller monomeric proteins). We first evaluated the iterative hybridization protocol (23) previously used to improve models generated using direct contacts predicted from GREMLIN on the entire set of CASP13 and CAMEO targets (Fig. S5). Incorporating our network derived distance predictions resulted in consistent improvement in model quality when the starting model’s TM-score was over 0.7, in a few cases by more than 10% in TM-score. We also tested the incorporation of the network derived distance restraints into the more CPU-intensive structure refinement protocol we used in CASP13 (10) on the CASP13 FM targets with an estimated starting TM-score > 0.6 that were not heavily intertwined oligomers and not bigger than 250 residues. Consistent improvements were observed on a set of six such targets (Fig. S6), with an average TM-score improvement of about 4%. The net improvement in prediction for these targets using the combination of the new structure generation method and refinement using the distance predictions is indicated by the red points in Fig. 3B.

### Assessing the ideality of *de novo* protein designs

Following up on the AlphaFold group’s excellent CASP13 prediction of the designed protein T1008, we systematically compared the ability of trRosetta to predict the structure of *de novo* designed proteins from single sequences compared to native proteins in the same length range. We collected a set of 18 *de novo* designed proteins of various topologies (24–26) (α, β and α/β) with coordinates in the PDB and a set of 79 natural proteins of similar size selected from the CAMEO set and ran trRosetta protocol to predict inter-residue geometries (Fig. 4A) and 3D models (Fig. 4B, examples of 3D models are in panels C-E). There is a clear difference in performance for natural proteins and *de novo* designs: the latter are considerably more accurate. The predicted structures of the designed proteins are nearly superimposable on the crystal structures, which is remarkable given that there is no co-evolution information whatsoever for these computed sequences, which are unrelated to any naturally occurring protein.

**Fig. 4.**
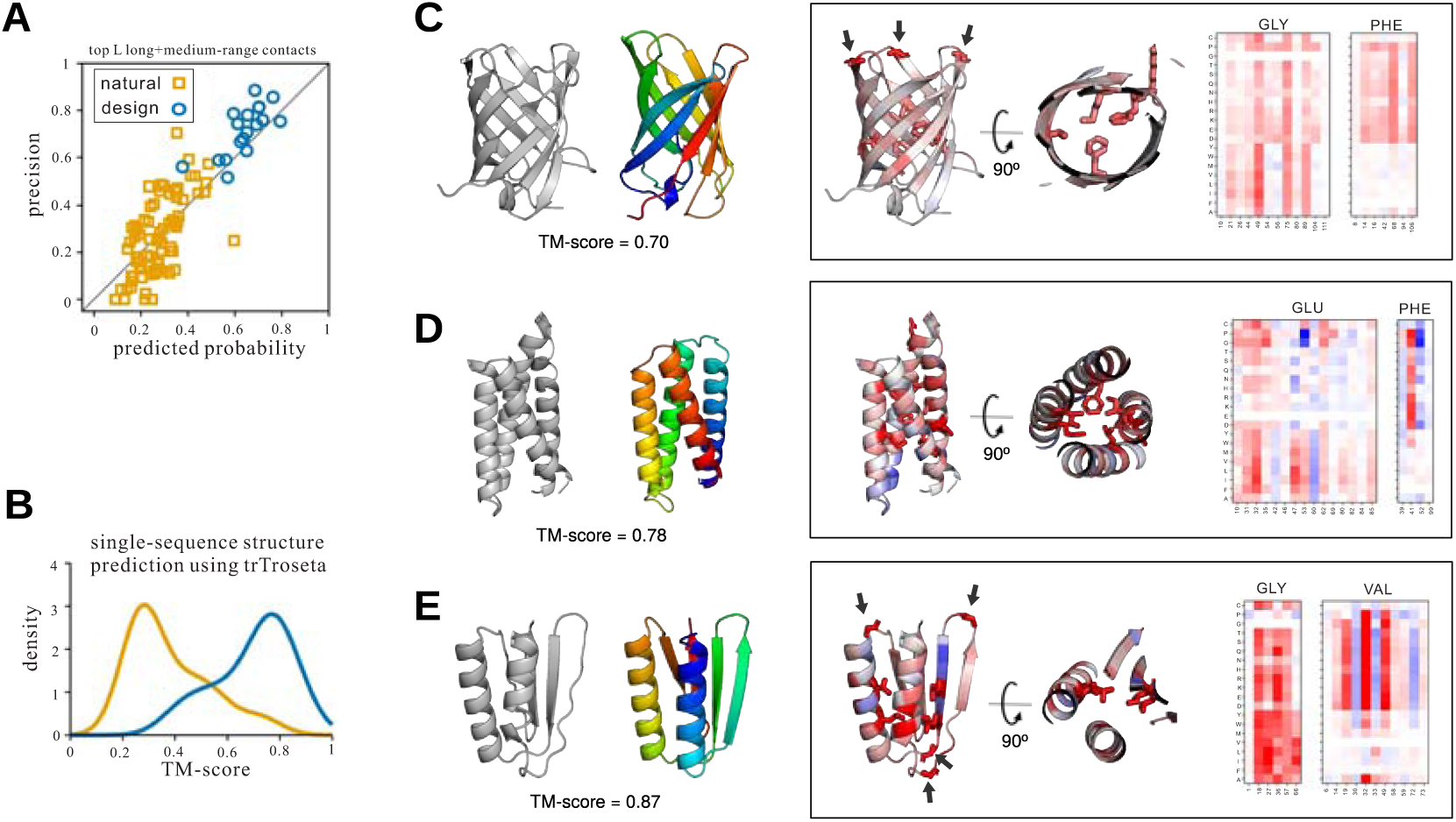
trRosetta accurately predicts structures of de novo designed proteins and captures effects of mutations. Differences in the accuracy of predicted contacts (A) and trRosetta models (B) for *de novo* designed and natural proteins of similar size from single amino acid sequences (C-E) Examples of trRosetta models for *de novo* designs of various topology: (C) non-functional β-barrel, PDB ID 6D0T, (D) α-helical IL2-mimetic, PDB ID 6DG6, (E) Foldit design with α/β topology, PDB ID 6MRS. Experimental structures are in gray and models are in rainbow. Framed panels show experimental structures color-coded by estimated tolerance to single-site mutations (red - less tolerant, blue - more tolerant); the 8 residues least tolerant to mutation are in stick representation, and glycine residues are indicated by arrows. Heat maps on the right show the change in probability of the designed fold for substitutions of the same residue type (indicated at top) at different sequence positions (indicated at bottom).

The high-accuracy structure prediction in the absence of co-evolutionary signal suggests the model is capturing fundamental features of protein sequence-structure relationships. To further investigate this, we performed an exhaustive mutational scanning of the ‘wild-type’ sequences for three designs of distinct topology (24–26) (Fig. 4C-E and Fig. S7). For each single amino acid substitution at each position, we calculated the change in the probability of the top *L* long+medium-range contacts (-log(*P*_*mutant*_/*P*_*WT*_). Mutations of core hydrophobic residues and of glycine residues in the β-turns produced large decreases in the probability of the designed structure. The effects of mutations depend strongly on context: the substitutions of the same amino acid type at different positions produce quite different changes in probability (Fig. 4C-E) which go far beyond the averaged out information provided by simple substitution matrices like BLOSUM or PAM.

## Discussion

The results presented here suggest that the orientation information predicted from co-evolution can improve structure prediction. Tests on the CASP13 and CAMEO sets suggest that our combined method outperforms all previously described methods, as it should as we have attempted to build on the many advances made by many groups in CASP13. However, it should be emphasized that retrospective analyses such as those carried out in this paper are no substitute for blind prediction experiments (as in the actual CASP13 and CAMEO), and that future CASP and CAMEO testing will be essential. Although not fully explored in this work, the integrated network architecture allows for backpropagation of gradients down to the MSA processing step, making it possible to learn optimal sequence reweighting and regularization parameters directly from data rather than using manually tuned values. To enable facile exploration of the ideas presented in this paper and in CASP13, the codes for the orientation prediction from co-evolution data, and the Rosetta protocol for structure generation from predicted distances and orientations are all available at http://yanglab.nankai.edu.cn/trRosetta/ and https://github.com/gjoni/trRosetta.

The accurate prediction of the structure of *de novo* designed proteins in the complete absence of co-evolutionary signal has implications for both the model and protein design generally. First, the model is clearly learning general features of protein structures. This is not surprising given that the direct couplings derived by the co-evolutionary analysis on a protein family are the two-body terms in a generative model for the sequences in the family, and thus training on these couplings for a large number of protein families is equivalent to training on large sets of protein sequences for each structure in the training set. From the design point of view, we have asserted previously that *de novo* designed proteins are “ideal” versions of naturally occurring proteins (27); the higher probability assigned by the model to designed proteins compared to naturally occurring proteins makes this assertion quantitative. Remarkably, similar “ideal” features appear to have been distilled from native protein analysis by expert protein designers to be incorporated into designed proteins, and extracted by deep learning in the absence of any expert intervention. Our finding that the model provides information on the contribution of each amino acid in a designed protein to the determination of the fold by the sequence suggests the model should be directly applicable to current challenges in *de novo* protein design.

This work also demonstrates the power of modern deep learning packages such as TensorFlow in making deep learning model development accessible to non-experts. The distance and orientation prediction method described here performs comparably or better than models previously developed by leading experts (of course we had the benefit of their experience), despite the relative lack of expertise with deep learning in our laboratory. These packages have now opened up deep learning to scientists generally-the challenge is more to identify appropriate problems, datasets and features than to formulate and train the models. The method developed here is immediately applicable to problems ranging from cryoEM model fitting to sequence generation and structure optimization for *de novo* protein design.

## Methods

### Benchmark datasets

#### Training set for the neural network

To train the neural network for the prediction of distance and orientation distributions, a training set consisting of 15,051 protein chains was collected from the PDB. First, we collected 94,962 X-ray entries with resolution ≤ 2.5□ (PDB snapshot as of May 1st 2018), then extracted all protein chains with at least 40 residues, and finally removed redundancy at 30% sequence identity cut-off, resulting in a set of 16,047 protein chains with the average length of 250 amino acids. All the corresponding primary sequences were then used as queries to collect MSAs using the iterative procedure described below. Only chains with at least 100 sequence homologs in the MSA were selected for the final training set.

#### Independent test sets

Two independent test sets are used to test our method. The first is the 31 FM domains (25 targets) from CASP13 (first target released on May 1st 2018). The second one is from the CAMEO experiment. We collected 131 CAMEO hard targets released between 2018.12.08 and 2019.06.01 - along with all the models submitted by public servers during this period. Note that for the CASP13 dataset, the full protein sequences rather than the domain sequences are used in all stages of our method to mimic the situation of the CASP experiments.

#### MSA generation and selection

The precision of predicted distance and orientation distribution usually depends on the availability of an MSA with ‘good’ quality. A deep MSA is usually preferable but not always better than a shallow MSA (see the examples provided in (3)). In this work, five alternative alignments are generated for each target. The first four are generated independently by searching the Uniclust30 database (version 2018_08) with HHblits (version 3.0.3) (28) with default parameters at four different *e*-value cutoffs: 1E-40, 1E-10, 1E-3, and 1. The last alignment was generated by several rounds of iterative HHblits searches with gradually relieved *e*-value cutoffs (1e-80, 1e-70, …, 1e-10, 1e-8, 1e-6, and 1e-4), followed by the *hmmsearch* (version 3.1b2) (29) against the metagenome sequence database (20) in case not enough sequences were collected at previous steps. The metagenome database includes about 7 billion protein sequences from the following resources: (i) JGI Metagenomes (7835 sets), Metatranscriptomes (2623 sets) and Eukaryotes (891 genomes); (ii) UniRef100; (iii) NCBI TSA (2616 sets); (iv) genomes manually collected from various genomic centers and online depositories (2815 genomes). To avoid attracting distant homologs at early stages and making alignment unnecessary deep, the search was stopped whenever either of the two criteria were met: at least 2000 sequences with 75% coverage or 5000 sequence with 50% coverage (both at 90% sequence identity cutoff) were collected. The final MSAs for the test datasets are available at http://yanglab.nankai.edu.cn/trRosetta/.

### Inter-residue geometries prediction by deep residual neural networks

#### Protein structure representation

In addition to the traditional inter-residue distance matrices, we also make use of orientation information to make the representation locally informative. For a residue pair (*i, j*), we introduce *ω* dihedral between C_*α*_, C_*β*_ of one residue and C_*β*_, C_*α*_ of the other, as well as two sets of spherical coordinates centered at each of the residues and pointing to the C_*β*_ atom of the other residue. These six coordinates (*d, ω, θ*_*ij*_, *φ*_*ij*_, *θ*_*ji*_, *φ*_*j*i_) are sufficient to fully define the relative orientation of two residues with respect to one another. Additionally, as it will be described below, any biasing energy term defined along these coordinates can be straightforwardly incorporated as restraints in Rosetta.

#### Input features

All the input features for the network are derived directly from the MSA, and are calculated on-the-fly. The 1D features include: (*i*) one-hot-encoded amino acid sequence of the query protein (20 feature channels), (*ii*) position-specific frequency matrix (21 features: 20 amino acids + 1 gap), and (*iii*) positional entropy (1 feature). These 1D features are tiled horizontally and vertically and then stacked together to yield 2×42=84 2D feature maps.

Additionally, we extract pair statistics from the MSA. It is represented by couplings derived from the inverse of the shrunk covariance matrix constructed from the input MSA. First we compute one-site and two-site frequency counts 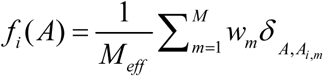 and 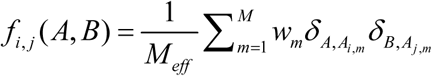, where *A* and *B*, denote amino acid identities (20 + gap), *δ* is the Kronecker delta, indices *i, j* run through columns in the alignment, and the summation is over all *M* sequence in the MSA; *w*_*m*_ is the inverse of the number of sequences in the MSA which share at least 80% sequence identity with sequence *m* (including itself); 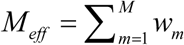. We then calculate the sample covariance matrix

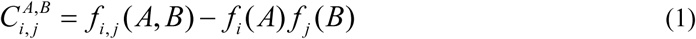

and find its inverse (also called the precision matrix) after shrinkage (i.e. regularization by putting additional constant weights on the diagonal):

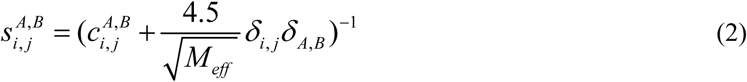

(More details on tuning the regularization weight in Eq. 2 are provided in Fig. S8). The 21×21 coupling matrices 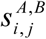 of the precision matrix (Eq. 2) are flattened, and the resulting *L*×*L*×441 feature matrix contributes to the input of the network. The above couplings (Eq. 2) are also converted into single values by computing their Frobenius norm for nongap entries:

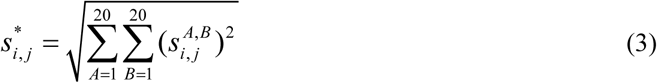

followed by the average product correction (APC):

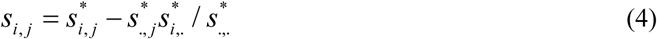

where 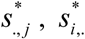 and 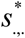 are row, column and full averages of the 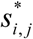 matrix, respectively. The coefficient in Eq. 2 was manually tuned on a non-redundant set of 1,000 proteins to maximize accuracy of the top *L* predicted contacts. From our experience, the final results are quite stable to the particular choice of the regularization coefficient in Eq. 2. To summarize, the input tensor has 526 feature channels: 84 (transformed 1D features) + 441 (couplings, Eq. 2) + 1 (APC score, Eq. 4).

#### Network architecture

The network takes the above *L*×*L*×526 tensor as the input, and applies a sequence of 2D convolutions to simultaneously predict 4 objectives: 1 distance histogram (*d* coordinate) and 3 angle histograms (*ω, θ* and *φ* coordinates). After the first layer, which transforms the number of input features down to 64 (2D convolution with filter size 1), the stack of 61 basic residual blocks with dilations are applied. Dilations cycle through 1, 2, 4, 8, 16 (12 full cycles in total). After the last residual block, the network branches out into 4 independent paths - one per objective - with each path consisting of a single 2D convolution followed by softmax activation. Since maps for *d* and *ω* coordinates are symmetric, we enforce symmetry in the network right before the corresponding two branches by adding transposed and untransposed feature maps from the previous layer. All convolution operations, except the first and the last, use 64 3×3 filters; ELU activations are applied throughout the network.

#### Training

We use categorical cross-entropy to measure the loss for all four objectives. The total loss is the sum over the four individual losses with equal weight (=1.0), assuming that all coordinates are equally important for structure modeling. During training, we randomly subsample the input MSAs, uniformly in the *log* scale of the alignment size. Big proteins of more than 300 amino acids long are randomly sliced to fit 300 residue limit. Each training epoch runs through the whole training set, and 100 epochs are performed in total. Adam optimizer with the learning rate 1e-4 is used. All trainable parameters are restrained by the *L*_2_ penalty with the 1e-4 weight. Dropout keeping probability 85% is used. We train five networks with random 95/5% training/validation splits, and use the average over the five networks as the final prediction. Training a single network takes ~9 days on one NVIDIA Titan RTX GPU.

### Structure determination by energy minimization with predicted restraints

#### Converting distance and orientation distribution to energy potential

The major steps for structure modeling from predicted distributions are shown in Fig. 1C. For each pair of residues, the predicted distributions are converted into energy potential following the idea of Dfire (30). For the distance distribution, the probability value for the last bin, i.e. (19.5, 20], is used as a reference state to convert the probability values into scores by the following equation.

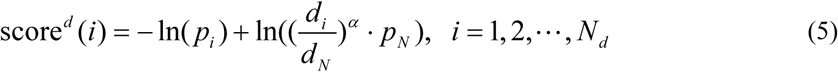

where *p*_*i*_ is the probability for the *i*-th distance bin, *N* is the total number of bins, *α* is a constant (=1.57) for distance-based normalization, *d*_*i*_ is the distance for the *i*-th distance bin. For the orientation distributions, the conversion is similar but without normalization, i.e.,

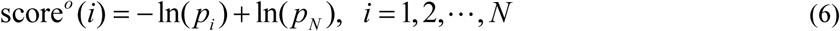

All scores are then converted into smooth energy potential by the *spline* function in Rosetta and used as restraints to guide the energy minimization. The range for distances is [0, 20Å] with a bin size of 0.5 Å, while for orientations, the ranges are [0, 360°] for *θ* and *ω*, and [0, 180°] for *φ*, all with a bin size of 15°; corresponding cubic spline curves are generated from the discrete scores defined by Eq. (5) or (6). For the distance-based potential, the *AtomPair* restraint is applied. For the *θ* and *ω*-based potential, the *Dihedral* restraint is applied. For the *φ*-based potential, the *Angle* restraint is applied.

#### Gradient-descent based energy minimization and full-atom based refinement

To speed up the modeling, coarse-grained (centroid) models are first built with the gradient-descent based energy minimization (*MinMover*) in Rosetta. A centroid model is a reduced representation of protein structure, in which the backbone remains fully atomic but each side chain is represented by a single artificial atom (CEN). The optimization is based on the L-BFGS algorithm (*lbfgs_armijo_nonmonotone*). A maximum of 1,000 iteration is used and the convergence cutoff is 0.0001. Besides the restraints introduced above, the following Rosetta energy terms are also used: ramachandran (rama), the omega and the steric repulsion van der Waals forces (vdw) and the centroid backbone hydrogen bonding (cen_hb). More details about these energy terms can be found in (16). The weights for the *AtomPair, Dihedral* and *Angle* restraints, rama, omega, vdw, and cen_hb are 5, 4, 4, 1, 0.5 and 1, respectively. The final models are selected based on the total score which includes both Rosetta energy and restraints scores.

The *MinMover* algorithm is deterministic but can be easily trapped into local minima. It is sensitive to the initial structure and restraints. Two strategies are proposed to introduce randomization effect and those models trapped into local minima can be discarded based on total energy. The first strategy is to use different starting structures with random backbone torsion angles (10 are tried). The second strategy consists in using different sets of restraints. For each residue pair, we only select a subset of restraints with probability higher than a specified threshold (from 0.05 to 0.5, with a step of 0.1).

For each starting structure, three different models are built by selecting different subsets of restraints based on sequence separation *s*: short range (1 ≤ *s* <12), medium range (12 ≤ *s* <24) and long range (*s* ≥ 24). The first one is progressively built with short-, medium- and long-range restraints. The second one is built with short+medium-range restraints and then with long-range restraints. The last one is built by using all restraints together.

In total, 150 (=10×5×3) centroid models were generated. The top 10 models (ranked by total energy) at each of the probability cutoff are selected for full-atom relax by *FastRelax* in Rosetta. In this relax, the restraints at probability threshold 0.15 are used together with the ref2015 scoring function. The weights for the *AtomPair, Dihedral* and *Angle* restraints are 4, 1 and 1, respectively.

## Data availability

The multiple sequence alignments for proteins in the benchmark datasets, the codes for inter-residue geometries prediction and the Rosetta protocol for restraint-guided structure generation are available at http://yanglab.nankai.edu.cn/trRosetta/ and https://github.com/gjoni/trRosetta.

## ACKNOWLEDGMENTS

We thank Frank DiMaio and David Kim for helpful discussions. This work was supported by National Natural Science Foundation of China (NSFC 11871290 to J.Y. and 61873185 to Z.P.), Fok Ying-Tong Education Foundation (161003, to J.Y.), KLMDASR (to J.Y.), Thousand Youth Talents Plan of China (to J.Y.), China Scholarship Council (to J.Y. and Z.P.), Fundamental Research Funds for the Central Universities (to J.Y.), National Institute of General Medical Sciences (grant no. R01-GM092802-07, to D.B.), National Institute of Allergy and Infectious Diseases (contract no. HHSN272201700059C, to D.B.), the Schmidt Family Foundation (to D.B.), and Office of the Director of the National Institutes of Health (grant no. DP5OD026389, to D.B. and DP5OD026389, to S.O.).

