## Supporting Information for "Improved protein structure prediction using predicted inter-residue orientations"

**Table S1.** The precision (%) of the derived contacts on the 31 CASP13 FM domains and the CAMEO targets compared with the top two contact predictors from CASP13. The values for other methods are slightly different from those listed on the CASP13 website, probably due to slightly different treatment of target length *L* (i.e., length of full sequence or length of domain structures; the latter is used here). The sequence separation between two residues *i* and *j* is denoted by *s* (=|*i*-*j*|).

| Method | $s \geq 24$ | | | $s \geq 12$ | | |
|---|---|---|---|---|---|---|
| | *L*/5 | *L*/2 | *L* | *L*/5 | *L*/2 | *L* |
| CASP13 FM domains | | | | | | |
| trRosetta | 78.5 | 66.9 | 51.9 | 89.9 | 83.1 | 70.2 |
| RaptorX-Contact | 70.2 | 57.8 | 44.7 | 87.3 | 76.1 | 61.3 |
| TripleRes | 65.5 | 54.8 | 42.3 | 84.7 | 74.6 | 60.9 |
| Baseline [a] | 68.6 | 56.5 | 44.3 | 80.0 | 71.6 | 60.7 |
| Baseline+*1* [b] | 70.7 | 59.8 | 46.0 | 83.9 | 74.3 | 62.2 |
| Baseline+*1+2* [c] | 74.0 | 61.8 | 48.2 | 84.6 | 77.3 | 64.6 |
| Baseline+*1+2+3* [d] | 78.1 | 65.1 | 51.3 | 89.9 | 82.8 | 69.3 |
| CAMEO hard / very hard targets | | | | | | |
| trRosetta | 82.7 / 75.4 | 72.0 / 63.2 | 57.0 / 48.0 | 82.7 / 81.5 | 76.4 / 74.8 | 64.4 / 62.8 |
| Baseline | 78.1 / 67.7 | 66.5 / 54.9 | 52.1 / 41.6 | 80.1 / 76.9 | 71.9 / 68.9 | 60.4 / 57.5 |
| Baseline+*1* | 79.5 / 71.6 | 67.3 / 57.7 | 52.5 / 43.1 | 81.8 / 78.8 | 73.3 / 69.8 | 60.5 / 57.4 |
| Baseline+*1+2* | 79.9 / 72.8 | 68.2 / 59.3 | 53.2 / 44.4 | 81.3 / 78.6 | 73.8 / 70.6 | 61.3 / 58.7 |
| Baseline+*1+2+3* | 80.0 / 73.1 | 70.0 / 60.9 | 54.9 / 46.1 | 81.5 / 80.5 | 74.9 / 73.4 | 62.8 / 61.4 |

[a] Baseline trRosetta model consists of 36 residual blocks and was trained without MSA subsampling or selection to predict distances only.
[b] *1* - adding MSA subsampling during training
[c] *2* - extending the network to predict orientations
[d] *3* - MSA selection based on predicted probability of the top *L* long+medium-range contacts

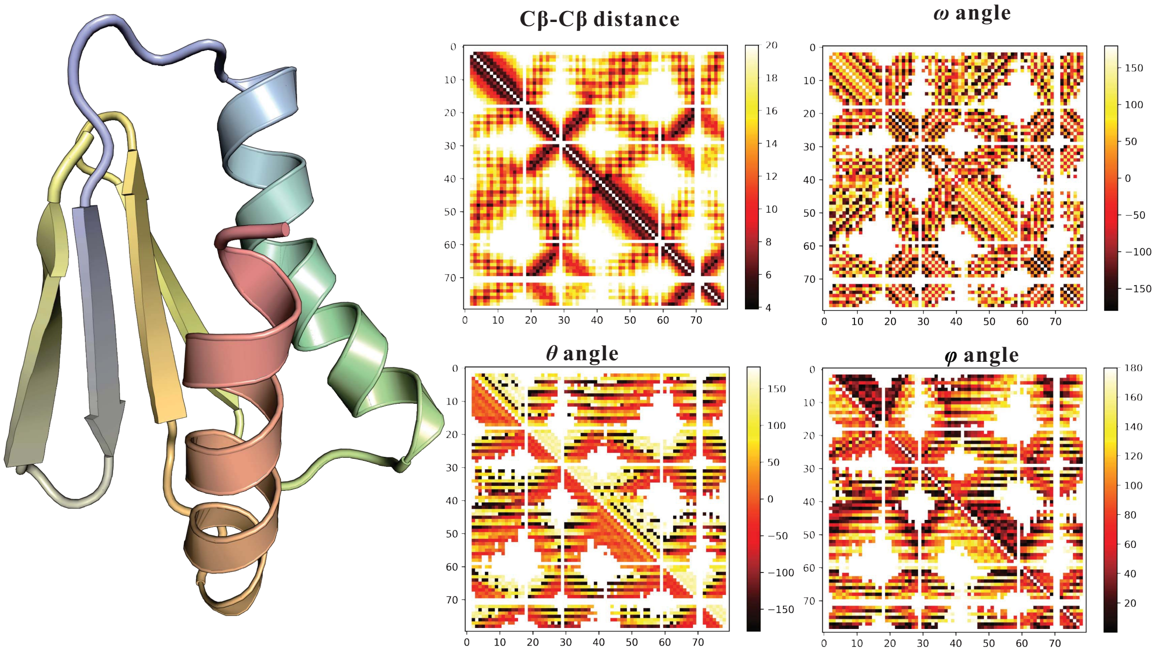


**Fig. S1**. *Visualization of inter-residue geometries for a representative structure (PDB ID: 6MSP)*. Colors on the 2D plots indicate inter-residue distances/angles.

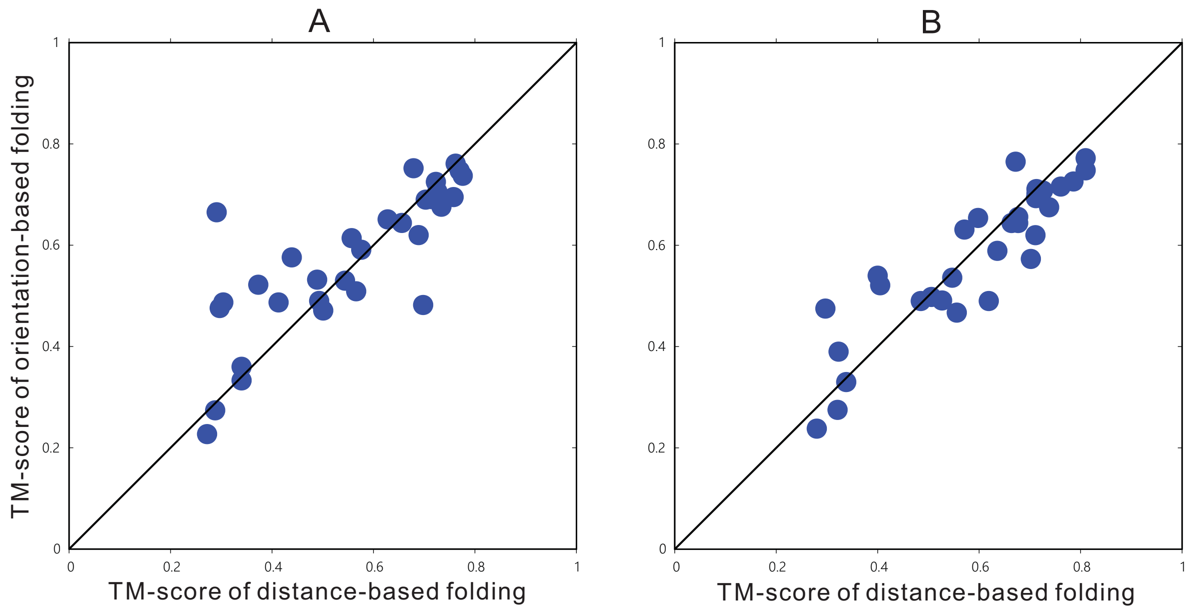


**Fig. S2**. *Comparison between models generated from distance and orientation-based folding.* (A) is for coarse-grained models and (B) is for relaxed models.

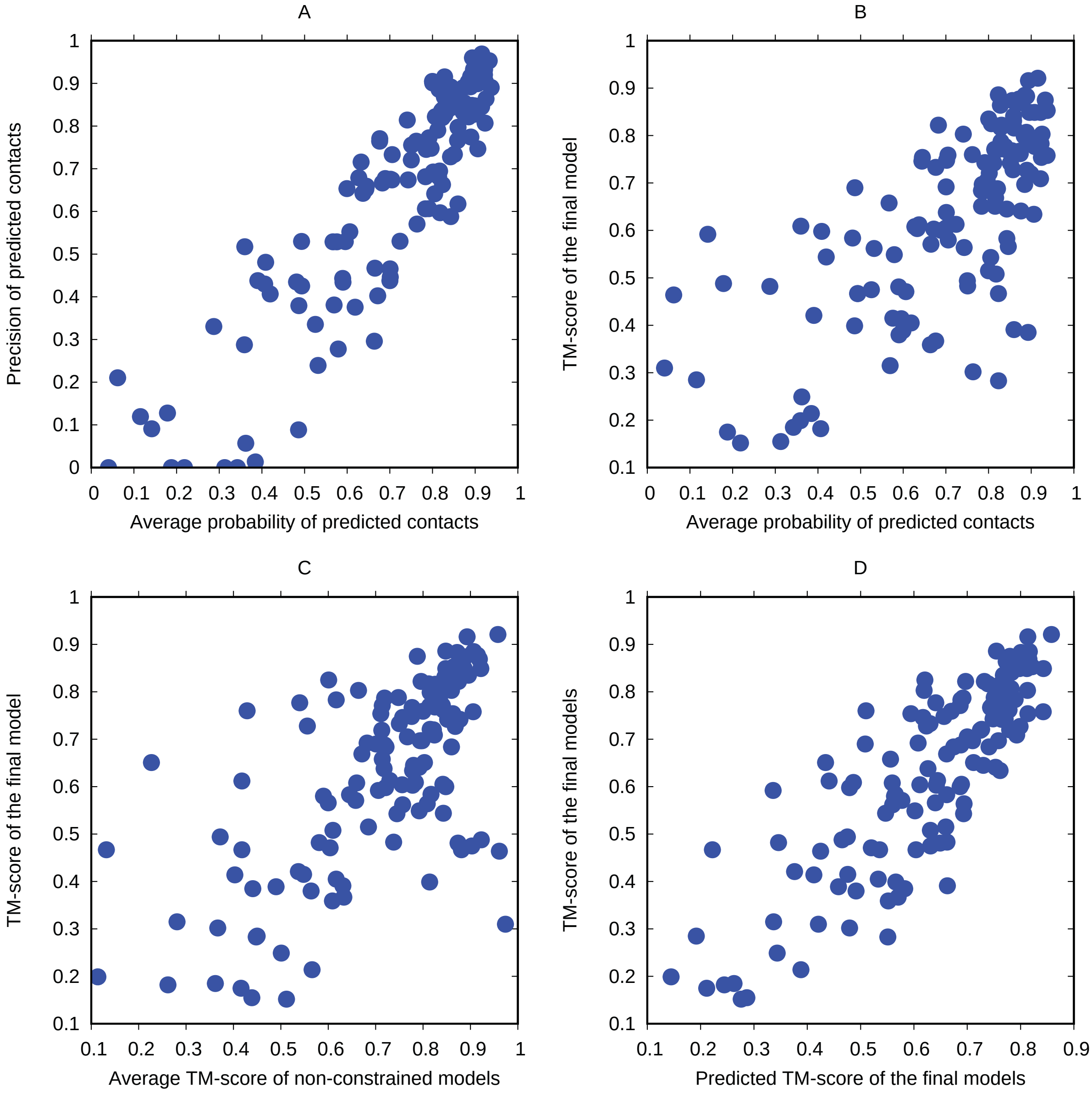


**Fig. S3**. *The correlation between different metrics on 131 CAMEO hard targets.*

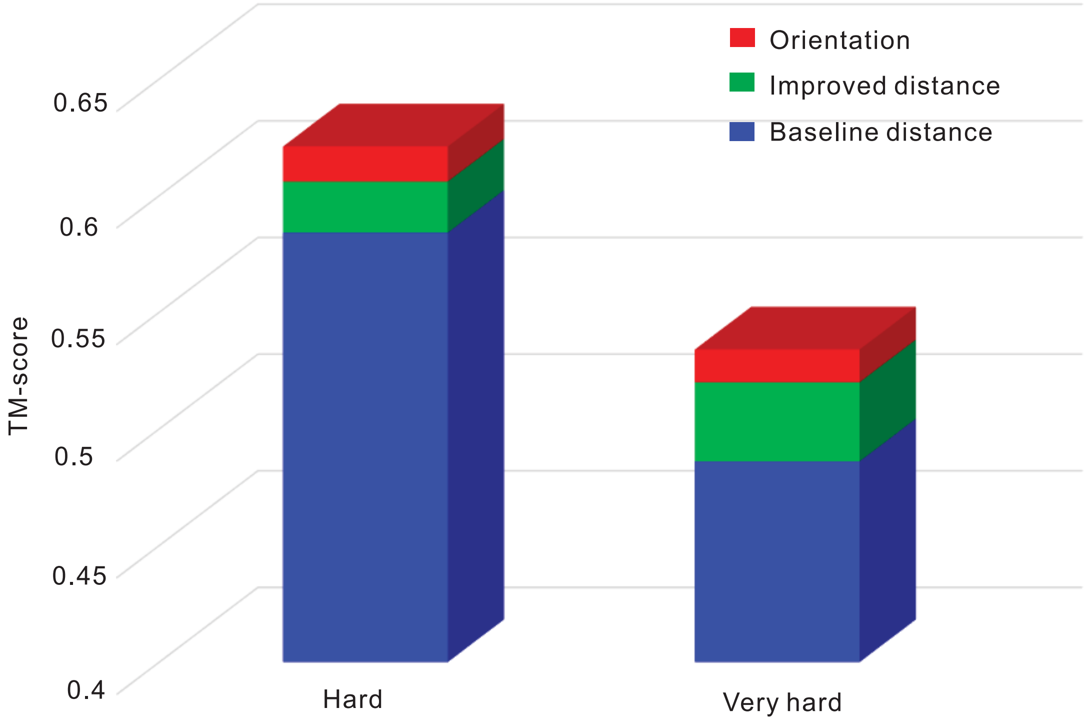


**Fig S4**. *Contributions of different components to our method on the CAMEO dataset.*

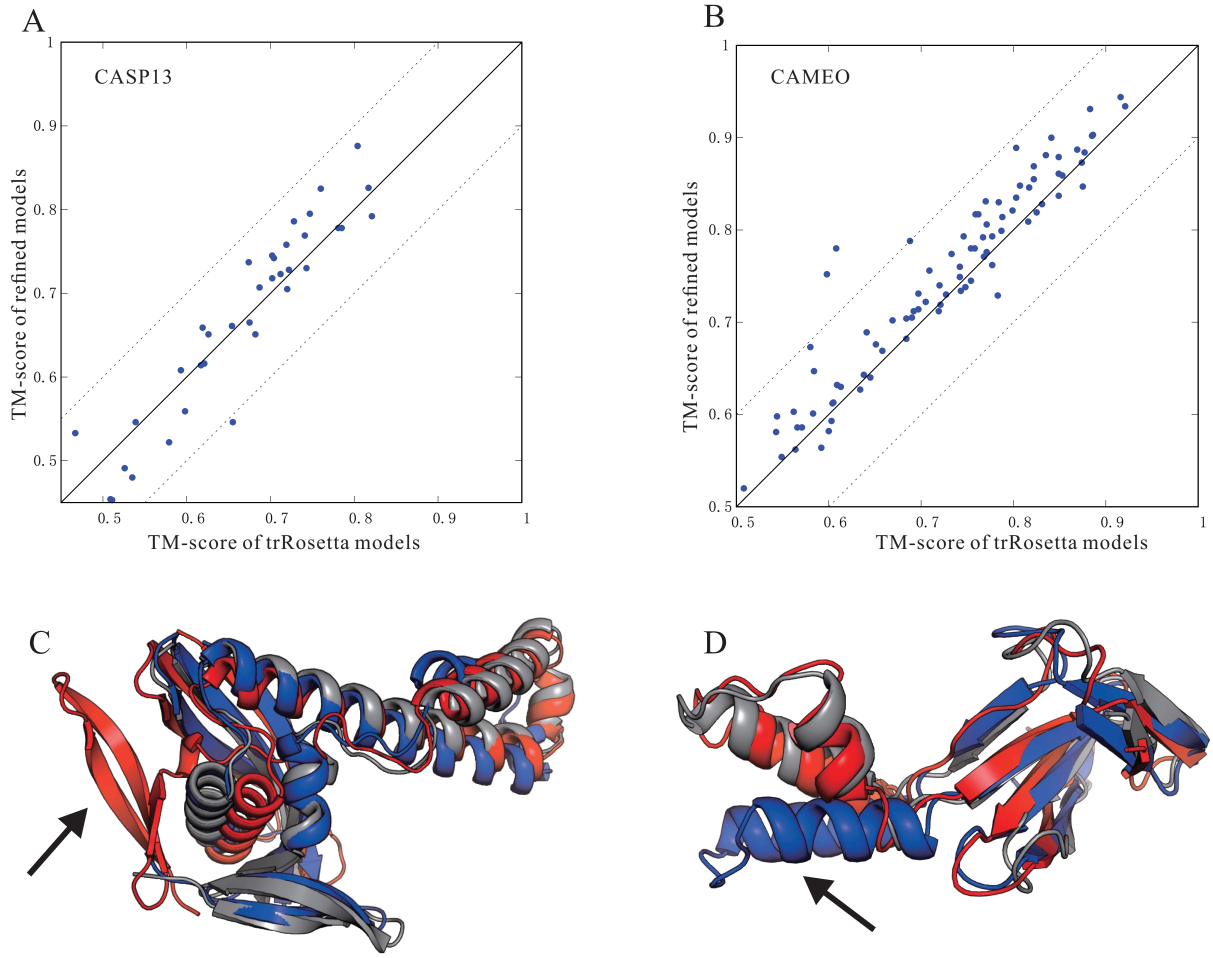


**Fig. S5**. *Refinement of models generated by trRosetta*. The best scoring model from trRosetta was further attempted for refinement with the protocol used in (Ovchinnikov et al, Science, 2017). Relative weight of 0.05 was applied to deep-learned distance restraints over Rosetta all-atom energy. Overall results for 31 CASP13 and 131 CAMEO targets are shown on panels (A) and (B) respectively. TM-score of trRosetta models and the corresponding relaxed models are shown in the x-axis and y-axis, respectively. (C, D) Representative examples of refinement results. Native, trRosetta model, and refined model are colored in gray, red, blue, respectively. (C) An example when refinement helped, 6CP8_B, improving TM-score from 0.61 to 0.78. (D) An example when refinement hurt, T1015s1, decreasing TM-score from 0.65 to 0.55.

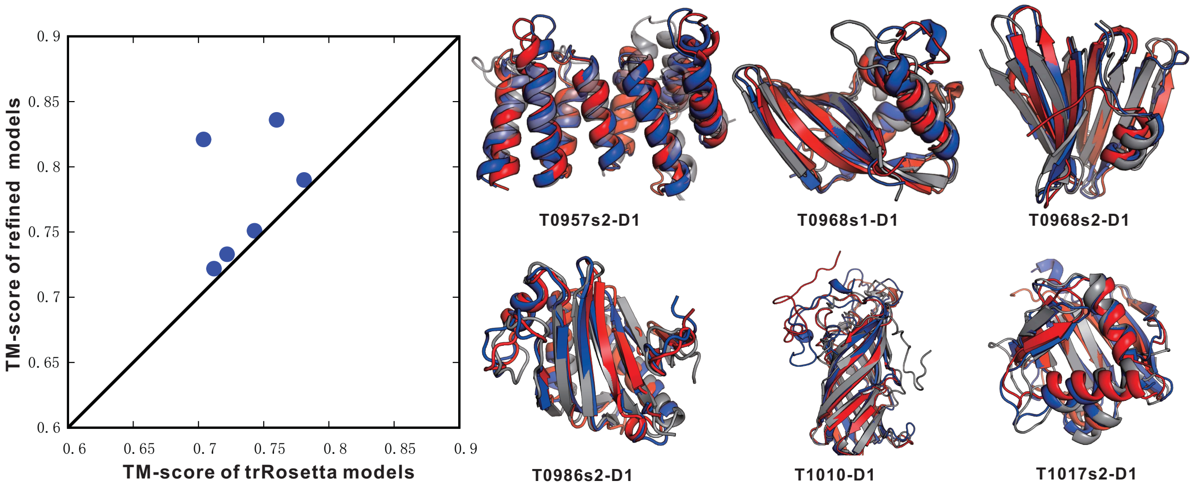


**Fig. S6**. *Accuracy of the refined models for six CASP13 targets by the protocol we used in CASP13.*

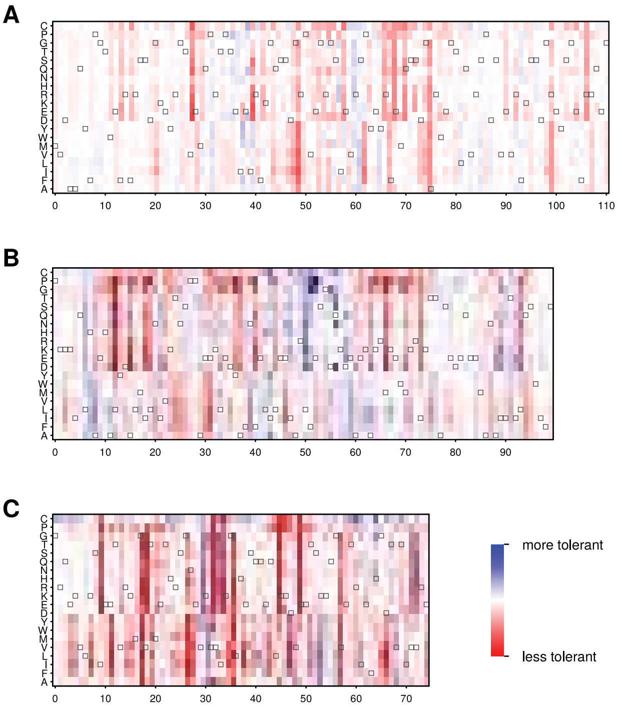


**Fig. S7**. *Effects of mutations as predicted by trRosetta for three de novo designed proteins of different topology*. (A) non-functional β-barrel, PDB ID 6D0T, (B) α-helical IL2-mimetic, PDB ID 6DG6, (C) Foldit design with α/β topology, PDB ID 6MRS (see Main text for details).

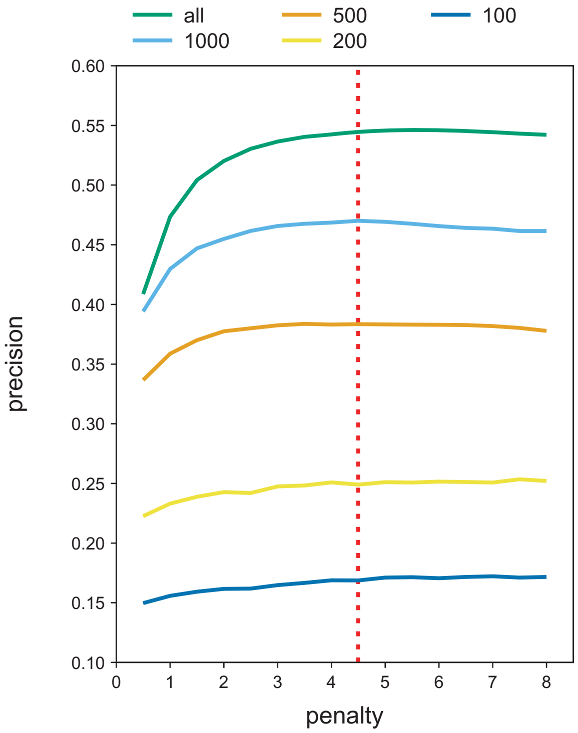


**Fig. S8**. *Effects of shrinkage strength on the precision of top L/2 long+medium-range contacts.* We systematically varied the penalty coefficient in Eq. 2 for alignments of different depth and selected the value (=4.5) which respects alignments at all diversity levels tested. We imposed the functional form of the shrinkage term to be *penalty**$M_{eff}^{-0.5}$ and only varied the *penalty* coefficient. No network was trained hereafter; the contacts were predicted directly from Eq. 4.
